# New strategy for bioplastic and exopolysaccharides production: Enrichment of field microbiomes with cyanobacteria

**DOI:** 10.1101/2023.05.30.542819

**Authors:** Beatriz Altamira-Algarra, Estel Rueda, Artai Lage, David San León, Juan F. Martínez-Blanch, Juan Nogales, Joan García, Eva Gonzalez-Flo

## Abstract

Seven photosynthethic microbiomes were collected from field environmental samples to test their potential in polyhydroxybutirate (PHB) and exopolysaccharides (EPS) production, two alternatives to chemical-based polymers. Microscope observations together with microbial sequence analysis revealed the microbiome enrichment in cyanobacteria after culture growth under phosphorus limitation. PHB and EPS production were studied under three culture factors (phototrophy, mixotrophy and heterotrophy) by evaluating and optimizing the effect of three parameters (organic and inorganic carbon and days under light:dark cycles) by Box-Behnken design. Results showed that optimal conditions for both biopolymers synthesis were microbiome-dependent; however, the addition of organic carbon boosted PHB production in all the tested microbiomes, producing up to 14%_dcw_ PHB with the addition of 1.2 g acetate·L^-1^ and seven days under light:dark photoperiods. The highest EPS production was 59 mg·L^-1^ with the addition of 1.2 g acetate·L^-1^ and four days under light:dark photoperiods. The methodology used in this article is suitable for enriching microbiomes in cyanobacteria, and for testing the best conditions for bioproducts synthesis for further scale up.

## Introduction

Interest in microbial biopolymers is growing due to increasing consumers’ concern about eco-friendly and sustainable products as an alternative to chemical-based polymers. Polyhydroxybutyrate (PHB) is a biodegradable polymer synthesized by numerous microorganisms as an energy and carbon storage product, while, exopolysaccharides (EPS) are excreted to protect the cells from environmental stresses or help them adhere to surfaces in the form of biofilms [1,2]. Both biopolymers have numerous applications in niche markets, such as textile, food, cosmetic, pharmaceutical and medical industries [1,3].

Cyanobacteria are a widespread group of photoautotrophic bacteria able to produce PHB and EPS using CO_2_ and solar energy. Consumption of CO_2_ for PHB and EPS production is a remarkable approach because it helps the mitigation of atmospheric greenhouse gases and contributes to the formation of a closed-carbon-loop for a polymer-based circular economy [4]. Nevertheless, cyanobacterial PHB and EPS industrial production and commercialization are still challenging due to their lower productivity compared to PHB synthesis by heterotrophic bacteria [5–8] or EPS production from plants and macroalgae [9,10]. In addition, current industrial production of both biopolymers results in high production costs due to the use of pure cultures, refined substrates and the need of sterile conditions [11,12], which hinders the introduction of these alternatives to the market.

PHB content in cyanobacteria and EPS release seems to rely on metabolism regime, in particular, production can be specially boosted by the addition of organic compounds to the medium. Mixotrophic regime (presence of organic and inorganic carbon sources) with acetate resulted in 46%_dcw_ PHB by *Anabaena* sp. after 7 days of incubation [13]. Interestingly, under heterotrophic conditions, *Synechocistys* sp. synthetized 22%_dcw_ PHB after 5 days in darkness and acetate supplementation [14]. Almost 30%_dcw_ PHB production was obtained by *Chlorogloea* sp. after 6 days in darkness and acetate as the only carbon source [15]. For EPS, a similar tendency is in principle predictable. However, studies regarding the effect of organic carbon on EPS production are very limited and clearly strain-dependent. For instance, *Arthrospira* sp. produced in mixotrophic condition 290 mg·L^-1^ in 4 days of incubation in presence of glucose and light, higher than production in photoautotrophy (220 mg·L^-1^) or heterotrophy (30 mg·L^-1^) alone.

In order to make these production processes a reality in an industrial context, microbiomes (microbial consortia) have emerged as a viable alternative to the pure-cultures. Microbiomes have the advantage to not need reactor sterilization, have a wider metabolic potential and extend feedstock options [16]. Current research focuses on biopolymer production by microbiomes, such as activated sludge from wastewater treatment plants, under heterotrophic regime using volatile fatty acids as carbon source [17–20]. However, a photosynthetic microbiome would combine the aforementioned advantages of working with a consortium, together with intrinsic benefits of photosynthetic microorganisms, which would use CO_2_ and light for growth and bioproducts synthesis. What we name as a photosynthetic microbiome may contain different types of microorganisms, including prokaryotic cyanobacteria, eukaryotic algae and non-photosynthetic microbes, such as heterotrophic bacteria. The principle behind the term “photosynthetic microbiomes” is that phototrophs are the dominant, but other microorganisms with different metabolisms can be simultaneously present. Nonetheless, exploration of biopolymer production by photosynthetic microbiomes is scarce. For example, [21] obtained a maximum concentration of 4.5%_dcw_ PHB by wastewater borne culture predominant in cyanobacteria. Also working with wastewater borne culture microbiome rich in cyanobacteria, [22] obtained 4.5%_dcw_ PHB using agricultural runoff as feedstock. Higher production was found by [23], who worked with a system enriched with proteobacteria and obtained 20%_dcw_ PHB content with acetate pulses.

In view of the above, in this work, different photosynthetic microbiomes were evaluated for the concurrent PHB and EPS production. These cultures were obtained from field environmental samples and enriched in cyanobacteria using low phosphorus (P) concentration as selective pressure. The aim of this study was to find whether microbiomes enriched in cyanobacteria produce PHB and EPS and identify the optimal metabolic regime (phototrophic, mixotrophic or heterotrophic) for biopolymer production. Metabolic regime was evaluated by combining three factors: (i) presence of organic carbon (OC), (ii) inorganic carbon (IC) and (iii) light:dark photoperiods. Multivariable experimental design (DoE) and surface response methodology (SRM) were used [24] to identify the optimal conditions and evaluate their effect (positive, negative or non) on PHB and EPS production to scale up the process in bigger photobioreactors.

In addition, this research combined microscope observations with advanced biomolecular techniques to validate the selective pressure applied to collected field environmental samples and identify communities in the photosynthetic microbiomes. This research was focused on evaluating conditions for PHB production but EPS were also analyzed to study the feasibility of coupling synthesis of both bioproducts. To the authors’ knowledge, this is the first time that microbiomes enriched with cyanobacteria are tested for PHB and EPS production under different regimes (photoautotrophic, mixotrophic and heterotrophic).

## Material and Methods

### Procurement of photosynthetic cultures

Field environmental samples were collected from (i) an urban pond located in an park (Barcelona, Spain, 41°24’31.0”N 2°12’49.9”E) fed with groundwater, (ii) Besòs river (Sant Adrìa de Besòs, Spain, 41°25’20.2”N 2°13’38.2”E), an intermittent Mediterranean river which receives high amounts of treated wastewater discharged from the wastewater treatment plants in the metropolitan area of Barcelona, (iii) Canyars canal outlet close to the sea (Gavà, Spain, 41°15’55.9”N 2°00’39.7”E), an area where water in the canal mixes with sea water, and (iv) the constructed wetland in Can Cabanyes (Granollers, Spain, 41°34’06.8”N 2°16’09.4”E), that receives treated water from the WWTP in Granollers (Table 1 and Fig. 1). Sample collection was done in two campaigns: (I) first campaign was performed in locations (i) and (ii) in June 2021; and (II) second campaign was done in locations (iii) and (iv) in October 2021.

**Table 1.** General characterization of samples.

| Campaign | Sample code | Origin of sample | pH | Conductivity<br>(mS·cm<sup>-1</sup>) | T<br>(°C) | N-NO<sup>3-</sup><br>(mg·L<sup>-1</sup>) | P-PO<sup>4-</sup><br>(mg·L<sup>-1</sup>) |
| --- | --- | --- | --- | --- | --- | --- | --- |
| First | UP | Urban pond | 8.04 | 1.7 | 26 | 0 | 0 |
|  | R1 | Besòs River |  |  |  |  |  |
|  | R2 | Besòs River | 8.03 | 1.7 | 23 | 3.8 | 0.35 |
|  | R3 | Besòs River |  |  |  |  |  |
| Second | CC | Canyars canal | 7.04 | 7.3 | 20 | 0.02 | 0 |
|  | CW1 | Constructed wetland | 7.6 | 1.5 | 20 | 0 | 0 |
|  | CW2 | Constructed wetland |  |  |  |  |  |

**Figure 1.**
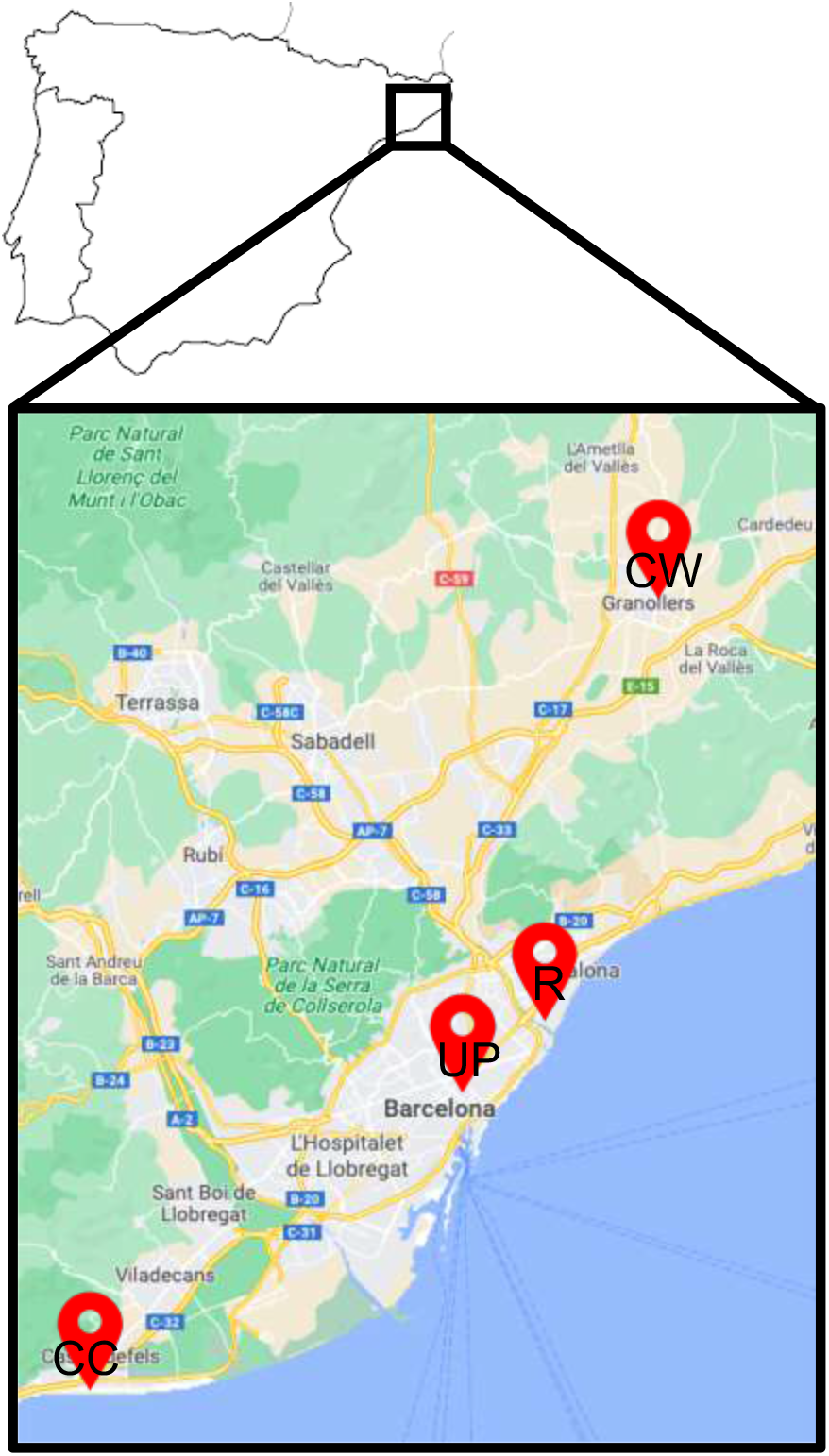
Location of sampling sites. CW: Constructed wetland; R: River, UP: Urban pond; CC: Canal

Samples were cultured in BG-11 medium with a low P concentration (0.2 mg·L^-1^) aiming to select cyanobacteria vs. other phototrophs. Cultures were grown under 5 klx illumination (HI 97500, HANNA instruments, Italy) (approx. 70 µmol m^-2^ s^-1^) in 15:9 h light:dark photoperiod provided by cool-white LED lights and continuously agitated by magnetic agitation. Biomass was scaled up to a larger volume every 15 days using a scale ratio of 1:5 (rueda2020) up to 1 L Erlenmeyer flasks. Microbiomes were maintained in these 1 L Erlenmeyer flasks by emptying 300 mL every week for biomass purge and subsequently adding 300 mL of fresh BG11 medium (up to a final low P concentration of 0.2 mg P·L^-1^). The selection and upscaling processes were daily monitored by bright light and fluorescence microscope (Eclipse E200, Nikon, Japan) observations. This biomass was the inoculum for the following tests (*Design of experiments*).

### Microbial identification

Cyanobacteria present in the microbiomes were identified and classified following morphological descriptions [25,26] under bright light microscope observation.

Molecular characterization was performed to identify the species by clone library based on 16S rRNA gene amplification carried out by the company ADM Biopolis. This analysis was performed on the four samples collected in the first campaign (UP, R1, R2, R3) six months after their collection and being under the mentioned selective pressure conditions. DNA was isolated following QIAmt Power fecal Pro DNA Kit (Werfen), adding bead beating and enzymatic lysis steps prior to extraction to avoid bias in DNA purification toward misrepresentation of Gram-positive bacteria. A total of 50 ng of DNA was amplified following the 16S Metagenomic Sequencing Library Illumina 15044223 B protocol (ILLUMINA).

In summary, in the first amplification step, primers were designed containing a universal linker sequence allowing amplicons for incorporation indexes and sequencing primers by Nextera XT Index kit (ILLUMINA); and 16S rRNA gene universal primers (Klindwoth2013). In the second and last assay amplification indexes were included. 16S based libraries were quantified by fluorimetry using QuantliT™ PicoGreen™ dsDNA Assay Kit (Thermofisher).

Libraries were pooled prior to sequencing on the MiSeq platform (Illumina), 250 cycles paired reads configuration. The size and quantity of the pool were assessed on the Bioanalyzer 2100 (Agilent) and with the Library Quantification Kit for Illumina (Kapa Biosciences), respectively. PhiX Control library (v3) (Illumina) was combined with the amplicon library (expected at 20%).

### Sequencing data analysis

Prior to sequence filtering and trimming, the sequences were tested for contaminations like chloroplasts. For this purpose, KRAKEN2 was used with the SILVA183 database without any type of filtering. Afterwards, the sequences were filtered by sequencing quality and Illumina primers were trimmed, and only sequences with identified primer are used for downstream analysis, in order to be able to carry out the amplicon reconstructions. PCR quimeras were filtered by QIIME2-DADA2.With these results a taxonomic annotation and abundance analysis was performed with nf-core/ampliseq v 2.2.0 [27] with Nextflow v20.05.0 using docker v20.10.8 using SILVA v183.1 database.

Downstream analysis was done with Microeco R package [Liu et al 2021].

### Design of Experiments

SRM based in Box-Behnken design was used to find the optimal condition for PHB and EPS production (phototrophic, mixotrophic or heterotrophic cultivation) and the microbiome with the highest biopolymer production. Three independent variables were evaluated: (i) acetate and (ii) bicarbonate concentrations, which were used as organic and inorganic carbon sources, respectively; and (iii) days in lightness. Three levels were assigned to each independent variable tested for each microbiome, selected according to previous results [24]. A total of 13 combinations (Table 2), including 3 replicates at the center point, were performed to fit the second order polynomial model, which allows determining the linear, quadratic and interaction effects of the studied factors in the PHB and EPS production [28]. The experimental design and the statistical analysis of results were done using the statistical software JMP (Cary. USA). A p-value<0.05 was applied as the significant level. SRM generates a second order polynomial expression that predicts the best response (highest concentration of PHB and EPS). This model was used to estimate the optimal cultivation conditions to maximize biopolymers production.

**Table 2.** Design matrix of Box-Behnken experiments.

| **Trial** | **Acetate (g C·L<sup>-1</sup>)** | **Bicarbonate (g C·L<sup>-1</sup>)** | **Days in light** | **Regime** |
| --- | --- | --- | --- | --- |
| 1 | 0 | 0 | 3.5 | - |
| 2 | 0 | 1 | 7 | Photoautotroph |
| 3 | 0 | 1 | 0 | Phototroph |
| 4 | 0 | 2 | 3.5 | Photoautotroph |
| 5 | 0.02 | 0 | 7 | Photoheterotroph |
| 6 | 0.02 | 0 | 0 | Heterotroph |
| 7* | 0.02 | 1 | 3.5 | Mixotroph |
| 8 | 0.02 | 2 | 7 | Mixotroph |
| 9 | 0.02 | 2 | 0 | Mixotroph |
| 10 | 0.04 | 0 | 3.5 | Photoheterotroph |
| 11 | 0.04 | 1 | 7 | Mixotroph |
| 12 | 0.04 | 1 | 0 | Mixotroph |
| 13 | 0.04 | 2 | 3.5 | Photoheterotroph |

\*this condition was done in triplicates, as determined by Box-Behnken design.

### 50 mL test tubes experiments

Conditions established by DoE were applied to all microbiomes. 50 mL Pyrex™ test tubes were inoculated with approximately 1 g VSS·L^-1^ and 50 mL BG11 medium without N and P in order to favour PHB accumulation [24]. Sodium acetate and sodium bicarbonate were added to the BG11 medium when required (Table 2). Tubes were continuously agitated by compressed sterile air bubbling and illuminated by cool-white LED lights, giving 2.1 klx (approx. 29 µmol m^-2^ s^-1^) in 15:9 h light:dark photoperiods. Dark condition was acquired by wrapping each tube in aluminium foil. Dissolved oxygen, pH, organic and inorganic carbon were measured at the beginning of the test. The cultivation conditions in Table 2 were maintained for 7 days. After, the aforementioned parameters together with PHB and EPS were analyzed as described in next section.

### Analytical methods

Biomass concentration was determined by analysis of total suspended solids (TSS) and volatile suspended solids (VSS) as described in Standard Methods 2540 C and 2540 D, respectively [29]. To assess photosynthetic activity and pH in the 50 mL tubes, dissolved oxygen and pH were measured offline by an oxygen meter (HI94142, HANNA Instruments) and pH meter (pH-meter, GLP 21, Crison, Spain), respectively.

To determine dissolved species, samples were previously filtered through a 0.7 μm pore glass microfiber filter. Nutrients (N and P) were measured as nitrate (N-NO_3_^-^) and phosphate (P-PO_4_^3^) following the methodology 4500-NO_3_^-^ and 4500-PE, respectively, described in Standard Methods [29]. Total and dissolved organic and inorganic carbon were analysed using a C/N analyser (2005, Analytikjena, Germany).

### PHB extraction and quantification

PHB analysis was adapted from methodology described in [30]. Briefly, 50 mL of mixed liquor were collected and centrifuged (4200 rpm, 7.5 min), frozen at −80 °C overnight in an ultra-freezer (Arctiko, Denmark) and finally freeze-dried for 24 h in a freeze dryer (−110 °C, 0.05 hPa) (Scanvac, Denmark). 3-3.5 mg of freeze-dried biomass were mixed with 1 mL CH_3_OH with H_2_SO_4_ (20% v/v) and 1 mL CHCl_3_ containing 0.05 % w/w benzoic acid. Samples were heated for 5 h at 100 °C in a dry-heat thermo-block (Selecta, Spain). Then, they were placed in a cold-water bath for 30 min to ensure they were cooled. After that, 1 mL of deionized water was added to the tubes and they were vortexed for 1 min. CHCl_3_ phase, containing PHB dissolved, was recovered with a glass pipette and introduced in a chromatography vial containing molecular sieves. Samples were analysed by gas chromatography (GC) (7820A, Agilent Technologies, USA) using a DB-WAX 125-7062 column (Agilent, USA). Helium was used as the gas carrier (4.5 mL min^-1^). Injector had a split ratio of 5:1 and a temperature of 230°C. FID had a temperature of 300°C. A standard curve of the co-polymer PHB-HV was used to quantify the PHB content.

### EPS composition analysis

15 mL of mixed liquor were centrifuged (4,400 rpm for 7.5 min). 4% NaCl and 96% cold ethanol in a proportion 2:1 (v/v) were added to the cell-free supernatant. Tubes were placed at 4°C overnight, followed by centrifugation (4,400 rpm, 20 min) in order to facilitate EPS precipitation. The pellet was freeze-dried (−110 °C, 0.05 hPa) (Scanvac, Denmark). Freeze-dried samples (∼5 mg) were dissolved in deionized water (5 mL) and hydrolyzed with 0.1 mL 99% trifluoroacetic acid (TFA) at 120°C for 4 h. The hydrolysate was used for the identification and quantification of the constituent sugar monomers and uronic acids by anion exchange chromatography, using a Metrosep Carb 2 -250/4.0 column (Metrohm, AG), equipped with a pulsed amperometric detector. The eluents used were (A) 1 mM sodium hydroxide and 1 mM sodium acetate, and (B) 300 mM sodium hydroxide and 500 mM sodium acetate. The analysis was performed at 30 °C, at a flow rate of 0.6 mL min^−1^.

## Results and Discussion

### Effect of selective pressure on field environmental samples

Microscope observations of samples UP, R1, R2 and R3, collected in the first campaign, revealed that initial environmental samples were very rich in microorganisms, such as green algae (e.g. *Scenedesmus* sp., *Cosmarium* sp. and *Chlorella* sp.), filamentous cyanobacteria (*Leptolyngbya* sp.) or diatoms (*Nitzschia* sp.) (Fig. 2). Biopolymers production strictly depends on culture composition; therefore, cultivation under P limitation was applied as a selective pressure to enrich them in cyanobacteria, as they have a higher capacity to store P intracellularly [21].

**Figure 2.**
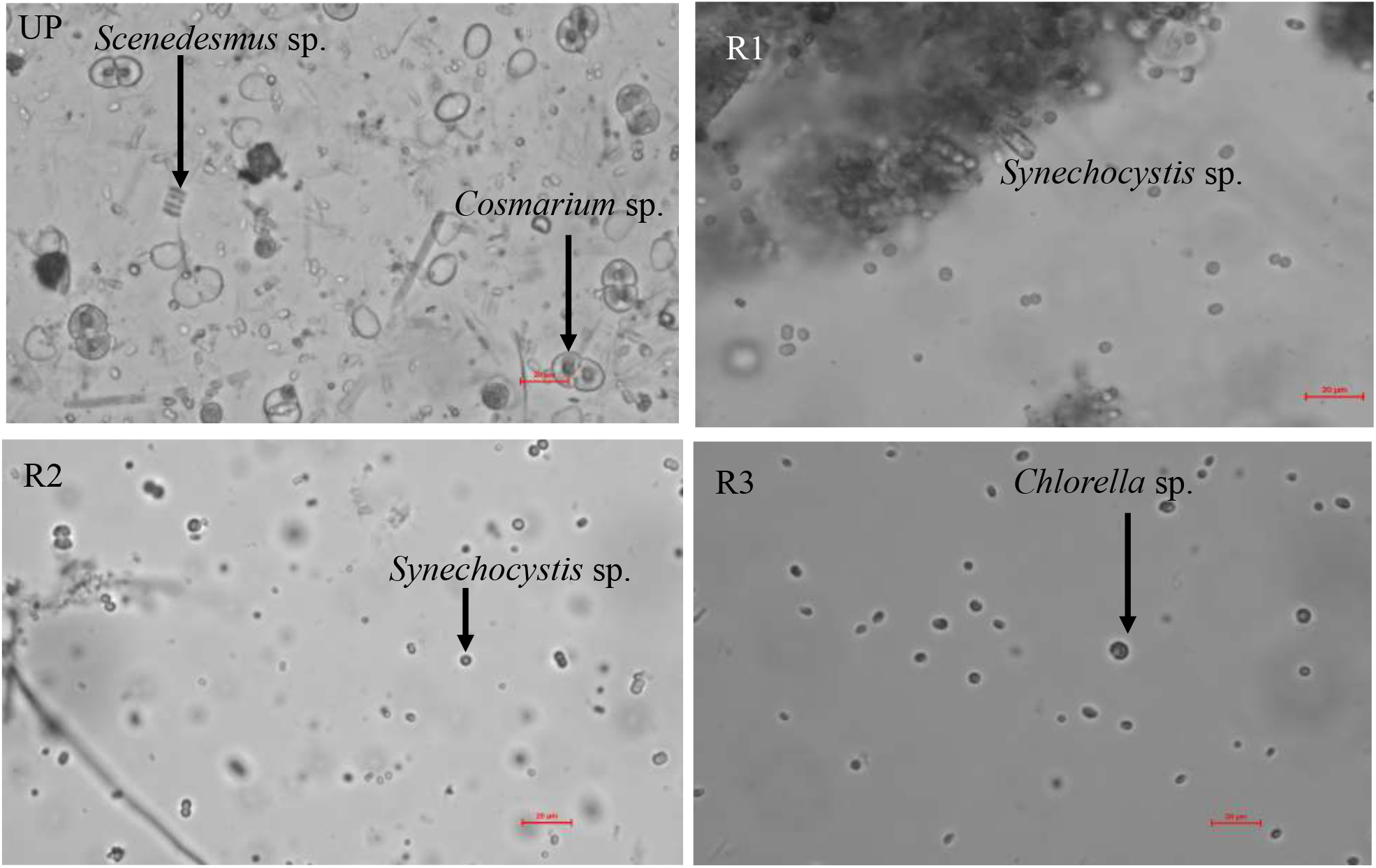
Microscope images of field microbiomes from UP, R1, R2 and R3 observed under bright light microscopy at 40X. Scale bar is 20µm.

Fig. 3 shows images of the microbiomes after 5 months of culture scale-up and growth under P limited conditions. It was clear that the P strict control enabled to enrich the microbiomes with cyanobacteria. Cyanobacteria species of *Leptolyngbya* sp., *Synechococcus* sp. and *Synechocystis* sp. were mostly observed. To validate microscope observations and identify bacterial species, highlthroughput 16S rRNA gene sequencing was applied to samples UP, R1, R2 and R3.

**Figure 3.**
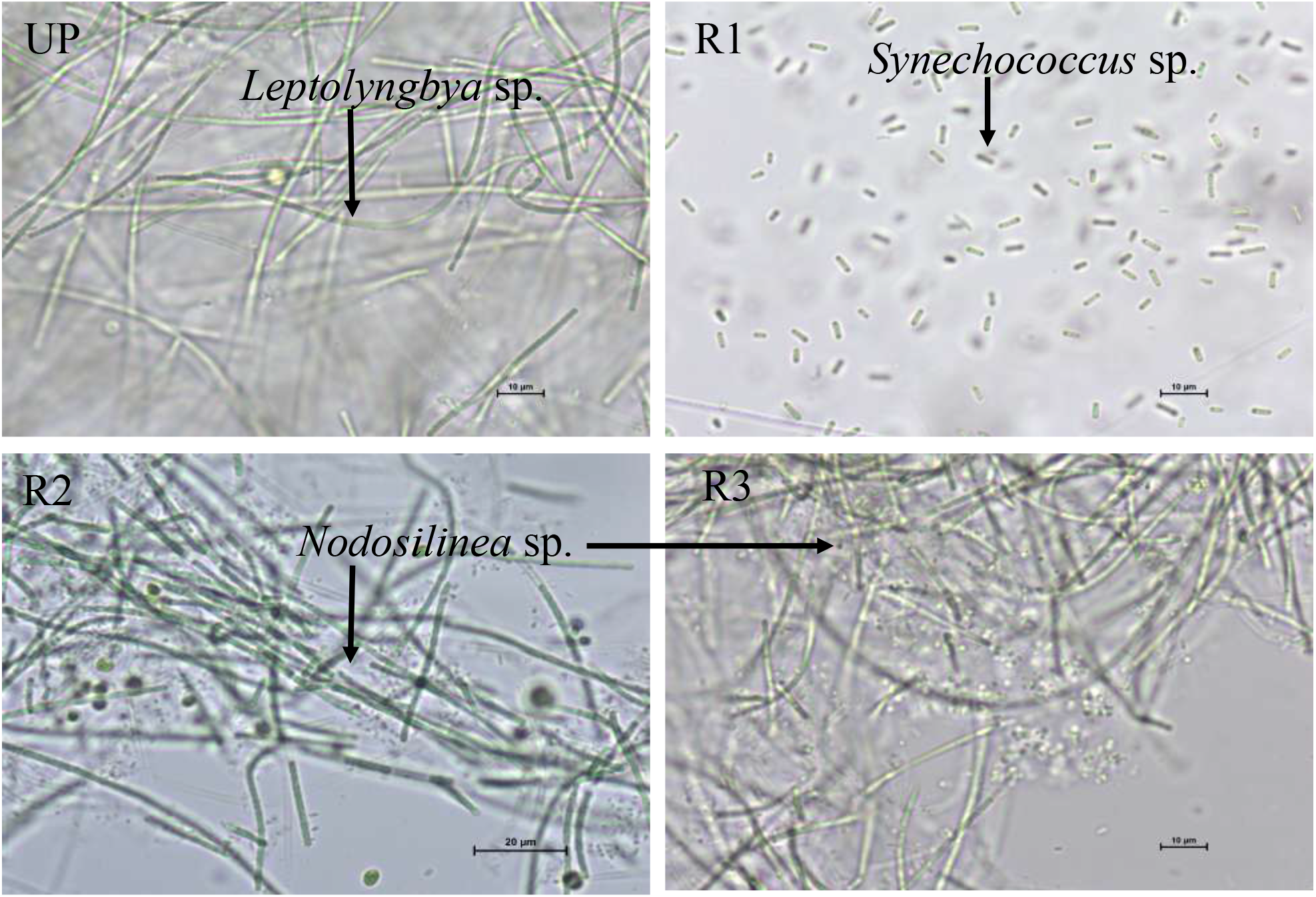
Microscope images of microbiomes from UP, R1, R2 and R3 observed under bright light microscopy at 40X after 5 months of growth with P limitation. Scale bar is 10 µm, except for R2, where scale bar is 20 µm.

### Microbial community analysis

The abundance of 16S sequences from selected samples confirm the high levels of cyanobacteria detected under the microscope. Other phyla like Bacteroidota and Proteobacteria were abundant too (Fig. 4). Nevertheless, there were differences in their relative frequency (RF) between the samples, mainly attributed to the different locations where initial samples were collected. For instance, 16s rRNA analysis revealed a microbial community in R1 mostly composed of Cyanobacteria (37%), as the RF of Bacteroidota and Proteobacteria (the second and third most abundant phyla) were 17% and 7%, respectively. High RF of photosynthetic microorganisms but similar to RF of other phyla were obtained in samples R3 (28% Cyanobacteria, 28% Bacteroidota and 31% Proteobacteria) and sample R2 (21% Cyanobacteria, 38% Bacteroidota and 21% Proteobacteria). In sample UP, RF of Cyanobacteria was the lowest of all the analyzed samples (12%).

**Figure 4.**
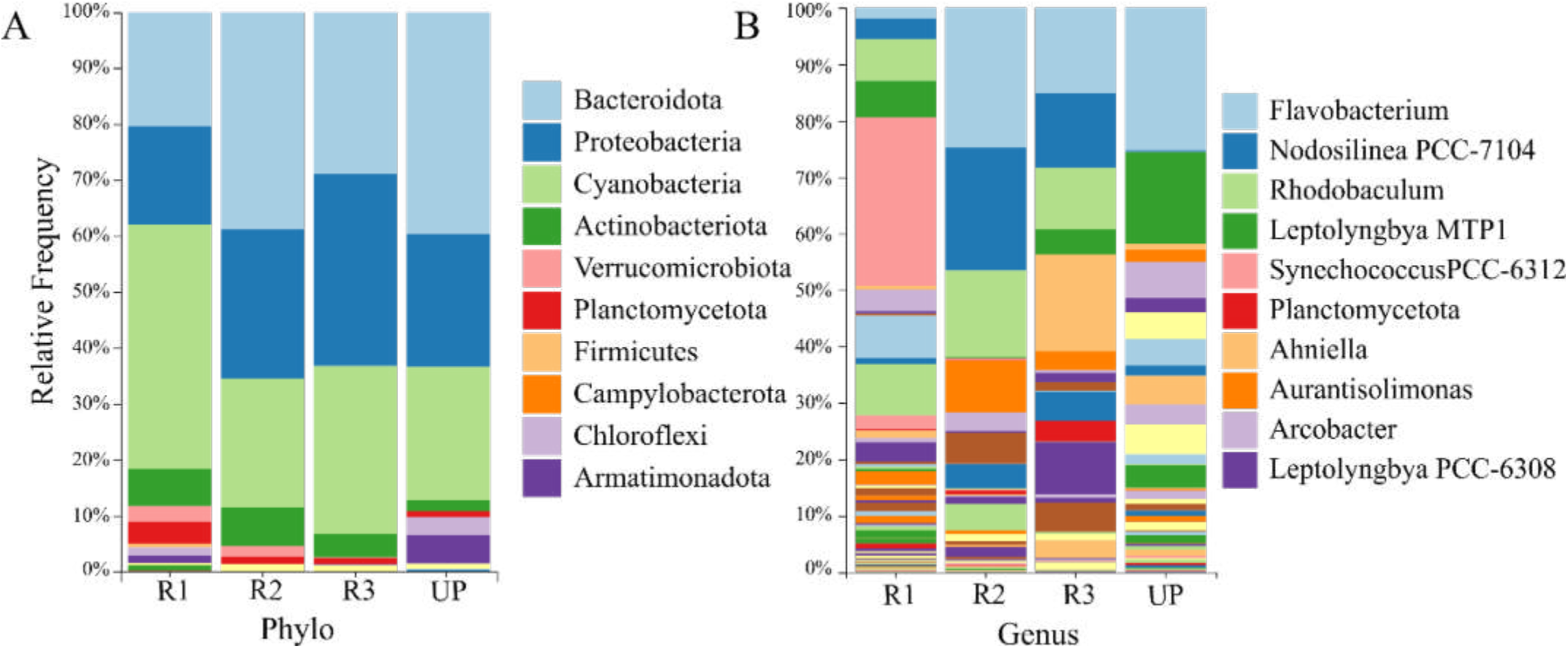
Relative abundance of microorganisms’ populations identified in the microbial community at (A) phyla level and (B) genus level after 5 months of growth with P limitation. The legend shows only the top 10 most abundant taxonomies.

Regarding the most abundant genus, cyanobacteria *Synechococcus sp*. was identified in sample R1 with 0.97 of confidence (Fig. A1) and a RF of 29%, being the most abundant bacteria with a notorious difference with respect to the RF of the second most abundant bacteria (9% *Cyclobacteriaceae*, a family of Bacteroidota). In samples R2 and R3, cyanobacteria *Nodosilinea* sp. was the second and third most abundant genera with a RF of 17% and 12%, respectively. *Leptolyngbya* sp. was the third genera most abundant in sample UP (6% of RF). In samples UP and R2, genus *Flavobacterium* (Bacteroidota) was the most abundant (up to 30% of RF).

As the selection pressure applied (P limitation) clearly enabled to obtain microbiomes rich in cyanobacteria, such process was further repeated in new collected environmental samples but without molecular analysis (samples CC, CW1 and CW2). Microscope observations were directly used to confirm the enrichment in cyanobacteria (Fig. 5). These samples were mostly enriched with the cyanobacterium *Synechococcus* sp. and they also have the presence of *Chlorella* sp.

**Figure 5.**
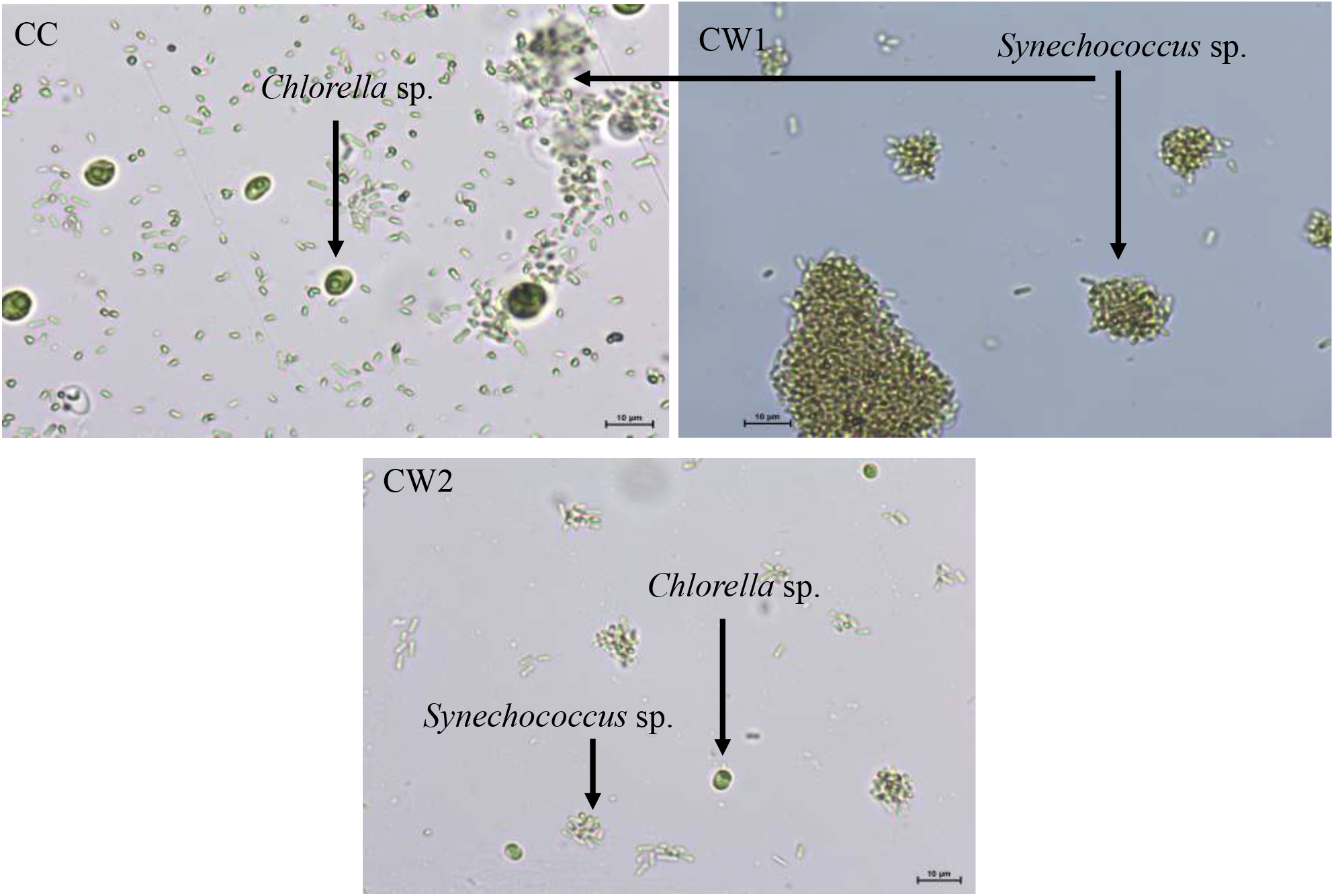
Microscope images of microbiomes from CC, CW1 and CW2 observed under bright light microscopy at 40X after 5 months of growth with P limitation. Scale bar is 10µm.

### PHB production under phototrophic, mixotrophic or heterotrophic regime

The effect of acetate (as the source of organic carbon, OC), bicarbonate (as the source of inorganic carbon, IC) and days in lightness on PHB production in samples UP, R1, R2, R3, CC, CW1 and CW2 was evaluated following a Box-Behnken design for RSM.

Acetate had a statistically significant (p-value<0.05) result on PHB production in all the microbiomes tested (Fig. A1). It triggered the synthesis in all microbiomes; however, it presented a quadratic and negative effect in samples CC, CW1 and CW2, meaning that concentrations higher than 0.8 g·L^-1^ would not contribute to increase PHB production (Table 3 and Fig. A1). For the other samples, acetate had a linear effect on PHB production (Fig. A1), meaning that biopolymer synthesis would be benefited from increasing OC concentration. Beneficial impact of OC on PHB accumulation has already been suggested for different pure cultures of cyanobacteria [14,15,24], which was attributed to the increase in the acetyl-CoA pool, which in turn activates the PHB metabolism [31]. Moreover, as shown in the prediction profilers in Fig. A1, acetate variations had the greatest effect on PHB production out of the three tested parameters.

**Table 3.** Optimal conditions predicted by SRM (second order polynomial expression) for PHB production.

| Microbiome | Acetate (g C·L<sup>-1</sup>) | Bicarbonate (g C·L<sup>-1</sup>) | Light (days) | PHB (%dcw) |
| --- | --- | --- | --- | --- |
| UP | 0.04 | 2 | 4 | 8 |
| R1 | 0.04 | 2 | 7 | 4 |
| R2 | 0.04 | 0 | 7 | 13 |
| R3 | 0.04 | 0 | 3 | 9 |
| CC | 0.03 | 0 | 7 | 14 |
| CW1 | 0.02 | 1.2 | 0 | 8 |
| CW2 | 0.03 | 1 | 7 | 8 |

Statistically significant differences of bicarbonate as IC source was observed for samples R3 and CW1. The effect was negative and linear for sample R3, meaning that the higher its concentration, the lowest the PHB production; but it was quadratic for CW1, which means that up to 1 g IC·L^-1^ would benefit biopolymer synthesis, but higher IC concentrations would decrease the production. For the rest of the samples, IC had no statistical effect (Fig. A1) therefore, in principle, there is no need to add IC for PHB production.

Days in lightness statistically affected PHB production in four microbiomes. A positive and linear effect was observed for samples R1 and CW2, meaning that accumulation under light would result in high production of PHB (Fig. A2. However, in samples UP and R3, more than 4 days of light:dark cycles reduced biopolymer accumulation (Table 3).

Altogether the DoE results suggest that higher PHB contents are achieved under heterotrophy or mixotrophy, as OC addition was the only factor affecting biopolymer production for all the microbiomes. IC and days in lightness, which supposedly would affect PHB synthesis under photoautotrophic regime, had a statistically significant impact in few specific microbiomes (Fig. A1).

According to [14,15,32] cyanobacteria can synthesize PHB under the different conditions tested in this work: phototrophic (use of IC source), mixotrophic (use of OC and IC) and heterotrophic (OC source supply) conditions. In the present study a maximum of 14 ± 1%_dcw_ of PHB was obtained in 7 days with the addition of 0.6 g acetate·L^-1^ under photoheterotrophic conditions (under light:dark periods and without IC) by microbiome CC (Fig. A3). This is in agreement with the optimal conditions predicted by DoE (Table 3): photoheterotrophic regime by the supplementation of 0.8 g acetate·L^-1^, seven days under light:dark cycles and no IC. Under these desirable conditions, DoE predicts a production of 14% _dcw_ of PHB. In addition, as can be seen in Fig. A2, cultures of CC under mixotrophic or heterotrophic regimes (conditions 5-12) produced similar concentration of PHB (8-14%_dcw_) indicating that microbiome CC is resilient to changes because it adapted to both regimes. Moreover, this result displays that IC had no effect on PHB synthesis (Fig. A1).

Experimental results also revealed that microbiome R2 produced 14 ± 1%_dcw_ of PHB under darkness cultivation, 0.04 g OC·L^-1^ and 1 g IC·L^-1^ (mixotrophic regime) (Fig. A1). Interestingly, DoE predicts a 13%_dcw_ of PHB under 7 days in light:dark cycles with the addition of 0.04 g OC·L^-1^ (Table 3).

No interactions between the tested parameters were observed, despite the interactions observed by [24] when working with monocultures of *Synechococcus* sp. and *Synechocystis* sp. [24] reported a negative interaction between (i) IC and days under light and (ii) OC and IC for *Synechococcus* sp. A positive interaction between days under light and OC was seen in *Synechocystis* sp.

### EPS production under phototrophic, mixotrophic or heterotrophic regime

Acetate supplementation had a statistically significant effect on EPS production in samples CC, CW1 and CW2 (Fig. A3). In samples CC and CW1 the effect was positive, meaning that the higher the OC concentration, the more EPS would be produced. However, the opposite trend occurred for CW2, where acetate addition caused a decrease in the polysaccharides content.

Interestingly, the addition of bicarbonate decreased EPS synthesis in all tested microbiomes; therefore, no IC is needed according to DoE predictions for optimal production (Table 4). However, in sample CW1, a positive interaction between OC and IC was observed (Fig. 6A): higher the concentration of both carbon sources, the higher the EPS production. In this microbiome carbon oversupplied was apparently metabolized and channeled to EPS production. This fact has already been observed in EPS synthesis in *Nostoc* sp. mixotrophic cultures [33].

**Table 4.** Optimal conditions predicted by SRM (second order polynomial expression) for EPS production.

| Microbiome | Acetate (g C·L<sup>-1</sup>) | Bicarbonate (g C·L<sup>-1</sup>) | Light (days) | EPS (mg ·L<sup>-1</sup>) |
| --- | --- | --- | --- | --- |
| UP | 0.04 | 0 | 7 | 8 |
| R1 | 0.04 | 0 | 0 | 7 |
| R2 | 0 | 0 | 0 | 29 |
| R3 | 0.04 | 0 | 7 | 7 |
| CC | 0.04 | 0 | 7 | 8 |
| CW1 | 0.04 | 0 | 7 | 57 |
| CW2 | 0.01 | 1 | 7 | 6 |

**Figure 6.**
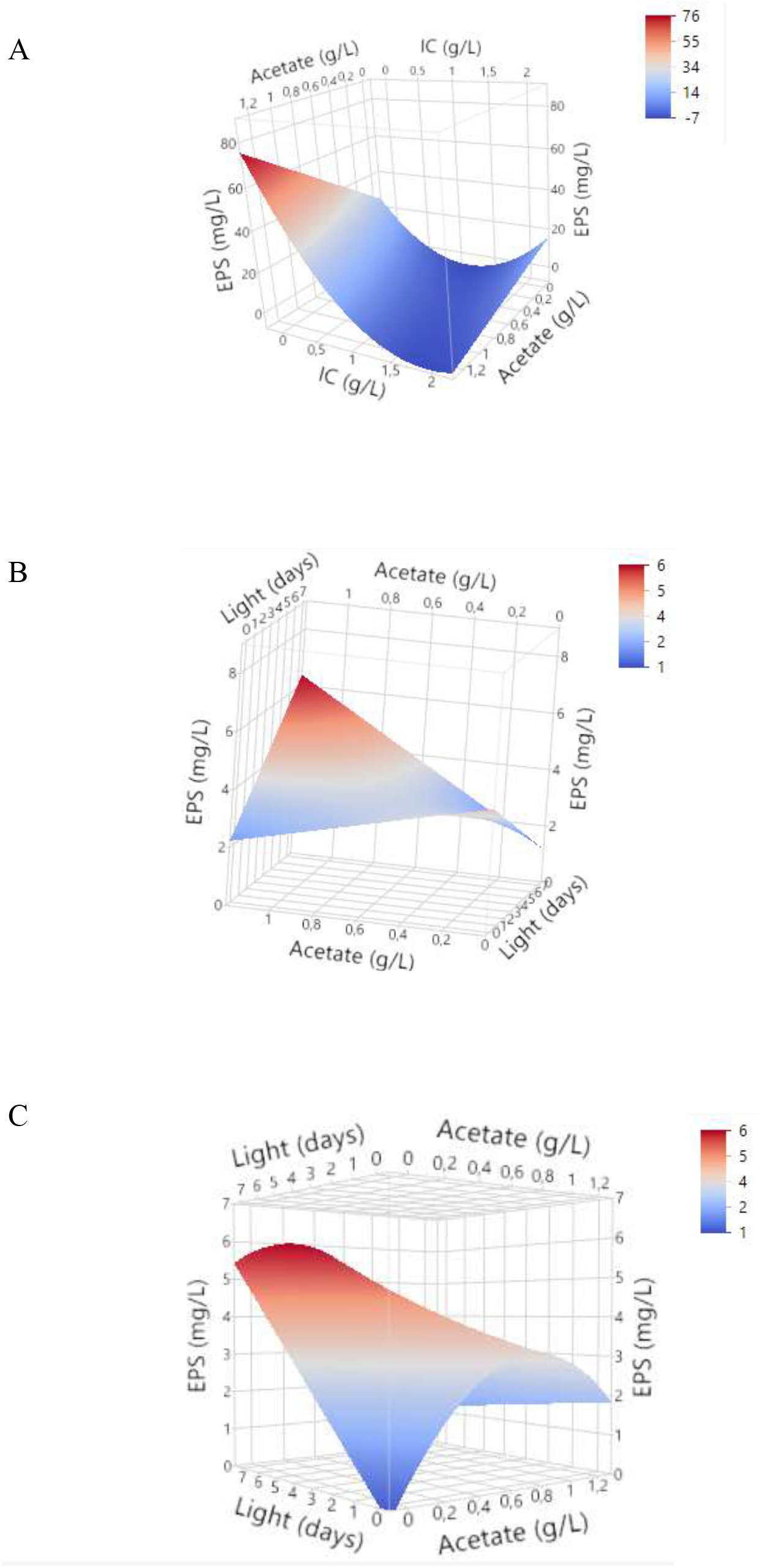
Surface plot of the interaction between (A) acetate and IC for microbiome CW1; (B) acetate and days under light:dark photoperiod for microbiome CC; and (C) acetate and days under light:dark photoperiod for microbiome CW2 on EPS production.

Significant interaction of acetate and light availability was found on EPS production in CC and CW2 (Fig. 6B and 6C). This interaction was different for each microbiome. In the case of CC this interaction was positive, so if the culture is kept under light:dark photoperiods, the increase in OC concentration will benefit EPS production. However, for CW2, the interaction was negative; when there was no OC addition, the more days under light, the higher the EPS production. This trend was also predicted by DoE as observed in Table 4.

The maximum EPS produced was 59 mg·L^-1^ by sample CW1 (Fig. A4) with the addition of 1.2 g acetate·L^-1^ and four days under light:dark photoperiods, which is similar to DoE predictions, where a maximum of 57 mg·L^-1^ could be produced under the supplementation of the same amount of acetate and seven days under light:dark photoperiods (Table 4). Still, this value is relatively low compared to EPS production in other cyanobacteria cultures. For example, [33] and [34] obtained 1,463 mg·L^-1^ and 634 mg·L^-1^, respectively. However, this higher EPS production was obtained in pure cultures of *Nostoc* sp. under mixotrophic regime and full nutrient availability. It is important to remember that here cultures were under N and P starvation; meaning that cell metabolism was restricted, to favor PHB synthesis. Therefore, it is hypothesized from our results that cells surviving under nutrient depletion, store the carbon sources present in the medium as carbon compounds rather than using it to produce EPS. Nonetheless, EPS productions seems to be strain dependent [35] and it is delicate to compare results by contrasting monocultures to microbiomes.

Regarding sugar composition of the EPS (Table A2), galactose and glucose were the most abundant monosaccharides produced in all samples and conditions out of the monosaccharides most frequently found in cyanobacterial EPS. Fucose and xylose were also produced by the microbiomes; however, production changed under cultivation conditions. Arabinose and mannose, which are also frequently produced by cyanobacteria, were barely present. Glucuronic was also present in all the samples. This uronic acid is also distinctive of cyanobacterial EPS.

Interestingly, optimal conditions for PHB and EPS seemed to be microbiome-dependent as results showed no clear relationship between taxonomic classification and biopolymers production. However, this hypothesis is not rejected. This event indicates that it is important to identify culture composition to stablish the operational process to maximize bioproduct synthesis. Here, microbiomes CC and CW2 could be used for PHB and EPS production as the optimal operation conditions for both biopolymers synthesis predicted by RSM coincided (Table 3 and 4), which would arise the possibility of coupling synthesis of both bioproducts for their compelling market entry. However, further research is necessary to obtain a highly functional microbiome and increase production yield.

## Conclusions

Phosphorus limitation successfully reduced bacteria and eukaryotic microalgae populations from field environmental samples and enabled to enrich the microbiomes cultures in cyanobacteria, as revealed by microscope observations and 16S rRNA analysis. Regarding to the operation conditions to produce PHB, acetate positively affected the biopolymer production in the seven tested microbiomes. The effect of inorganic carbon and days under light depended on the microbiomes, but the effect was clearly much lower in comparison to the effect of acetate. Parameters affecting EPS synthesis and production also depend on each microbiome, meaning that variables influencing biopolymer production depend on inoculum and culture composition. Interestingly, acetate supplementation boosted biopolymers production for almost all the microbiomes tested, suggesting a combine culture regime where biomass growth can be done in photoautotrophy (use of CO_2_ and light) and bioproducts synthesis in mixo-or heterotrophy regime (addition of OC and light, with or without IC). These results disclosed the potential of scaling up PHB production to bigger photobioreactors using R2 or CC as biomass, as well as, producing EPS by microbiome CW1. Results showed the feasibility of bioproducts synthesis with microbiome CC or CW1 but further research is needed to get a highly productive culture.

## CRediT authorship contribution statement

**Beatriz Altamira-Algarra:** Conceptualization, Validation, Formal analysis, Investigation, Writing – original draft. **Estel Rueda:** Conceptualization, Writing – review & editing. **Artai Lage:** Investigation. **David San León:** Sequencing data analysis, Writing – review & editing. **Juan F. Martínez-Blanch:** Genome sequencing. **Juan Nogales:** Sequencing data analysis, Writing – review & editing. **Joan Garcia:** Conceptualization, Resources, Writing – review & editing, Supervision, Project administration, Funding acquisition. **Eva Gonzalez-Flo:** Conceptualization, Supervision, Writing – review & editing.

## Appendix

A. Supporting information

Supplementary data associated with this article can be found in the online version at

