## Supplementary material for "New strategy for bioplastic and exopolysaccharides production: Enrichment of field microbiomes with cyanobacteria"

^a^ GEMMA-Group of Environmental Engineering and Microbiology. Department of Civil and Environmental Engineering. Escola d’Enginyeria de Barcelona Est (EEBE). Universitat Politècnica de Catalunya-BarcelonaTech. Av. Eduard Maristany 16. Building C5.1. E-08019 Barcelona. Spain

^b^ Department of Systems Biology, Centro Nacional de Biotecnología, CSIC, Madrid, Spain

^c^ Interdisciplinary Platform for Sustainable Plastics towards a Circular Economy-Spanish National Research Council (SusPlast-CSIC), Madrid, Spain

^d^ Department of preventive medicine, public health, food sciences, toxicology and forensic medicine, Universitat de Valencia, Valencia, Spain

^e^ Biopolis S.L., ADM, Parc Cientifc Universidad De Valencia, Edif. 2, C/ Catedrático Agustín Escardino Benlloch, 9, 46980 Paterna, Spain

^f^ GEMMA-Group of Environmental Engineering and Microbiology. Department of Civil and Environmental Engineering. Universitat Politècnica de Catalunya-BarcelonaTech. c/ Jordi Girona 1-3. Building D1. E-08034 Barcelona. Spain

**Table A1.** Relative frequency (%) of microorganisms from taxonomical classification (only shown those microorganisms with ≥ 5% of relative frequency in at least one sample).

|  |  |  |  |  |  | **Sample** | | | |  |
| --- | --- | --- | --- | --- | --- | --- | --- | --- | --- | --- |
| **Kingdom** | **Phylum** | **Class** | **Order** | **Family** | **Genus** | **UP** | **R1** | **R2** | **R3** | **Confidence** |
| Bacteria | Bacteroidota | Bacteroidia | Cytophagales | Cyclobacteriaceae |  |  | 9.04 |  |  | 0.88 |
|  |  |  | Flavobacteriales | Flavobacteriaceae | Flavobacterium | 38.01 |  | 24.23 | 14.71 | 0.62 |
|  |  |  | Chitinophagales | Chitinophagaceae | Aurantisolimonas |  |  | 9.20 | 3.25 | 1 |
|  |  |  | Sphingobacteriales | NS11-12 marine group |  | 4.49 |  | 4.21 | 5.07 | 1 |
|  |  |  | Cytophagales |  |  |  |  |  | 5.20 | 0.61 |
|  | Cyanobacteria | Cyanobacteriia | Leptolyngbyales | Leptolyngbyaceae | Leptolyngbya MTP1 | 1.67 | 5.45 |  | 4.44 | 0.85 |
|  |  |  | Leptolyngbyales | Leptolyngbyaceae | Leptolyngbya PCC-6306 |  |  |  | 8.80 | 1 |
|  |  |  | Phormidesmiales | Nodosilineaceae | Nodosilinea PCC-7104 | 1.35 | 2.67 | 21.37 | 12.01 | 1 |
|  |  |  | Thermosynechococcales | Thermosynechococcaceae | Synechococcus PCC-6312 |  | 29.22 |  |  | 0.97 |
|  | Proteobacteria | Alphaproteobacteria | Rhodobacterales | Rhodobacteraceae | Rhodobaculum | 7.45 | 6.585 | 15.25 | 10.87 | 1 |
|  |  |  | Acetobacterales | Acetobacteraceae | Roseococcus | 6.525 |  | 5.51 | 1.52 | 1 |
|  |  |  | Rhodobacterales | Rhodobacteraceae |  | 3.30 |  |  | 1.49 | 0.8 |

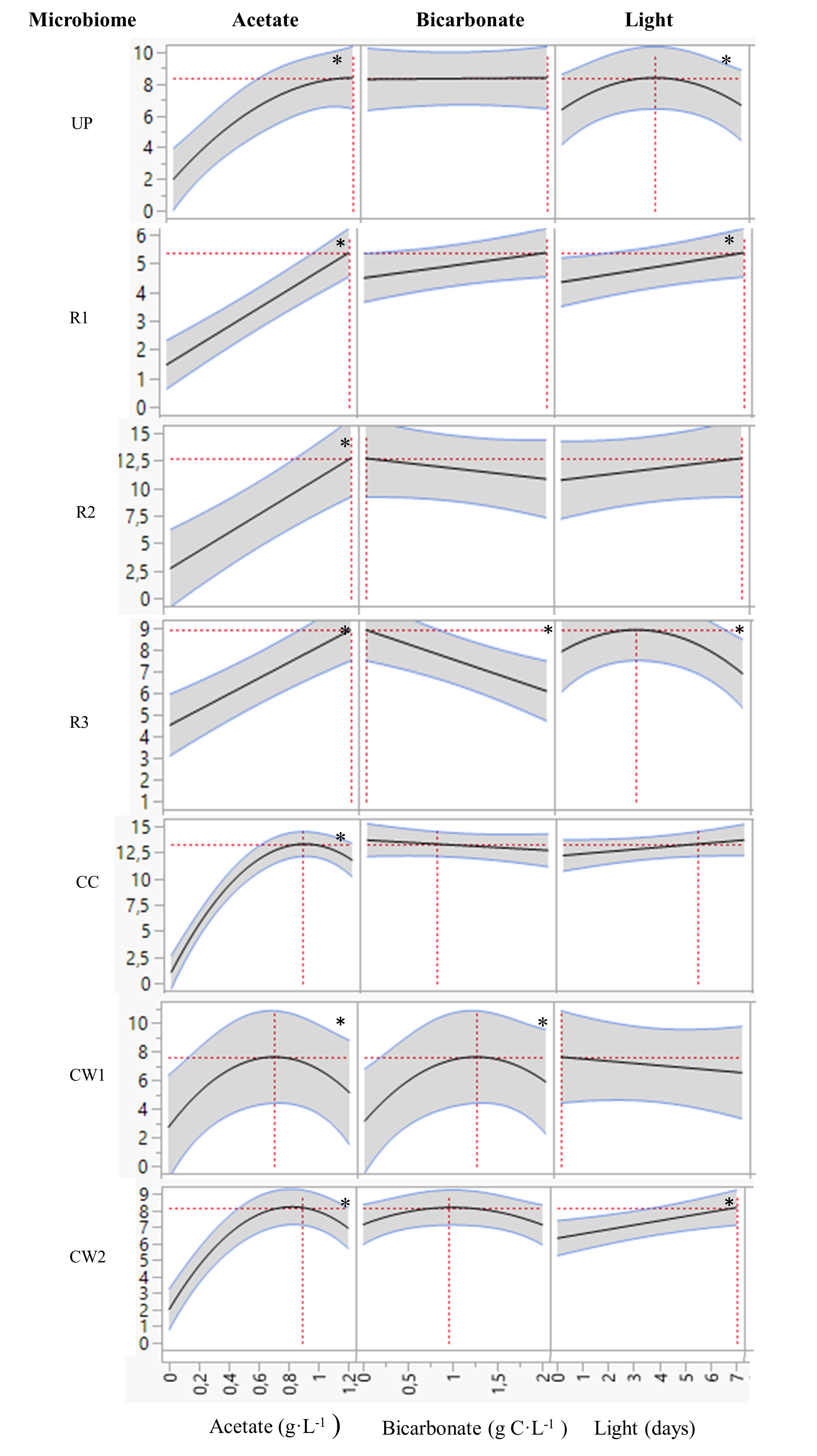

**Figure A1.** Prediction of the effect of acetate, inorganic carbon and light in the PHB production based on DoE results. Illustrating the optimal values of the tested parameters for each microbiome. * indicates parameters which effect is statistically significant. Regression line is shown in black and confidence interval in grey.

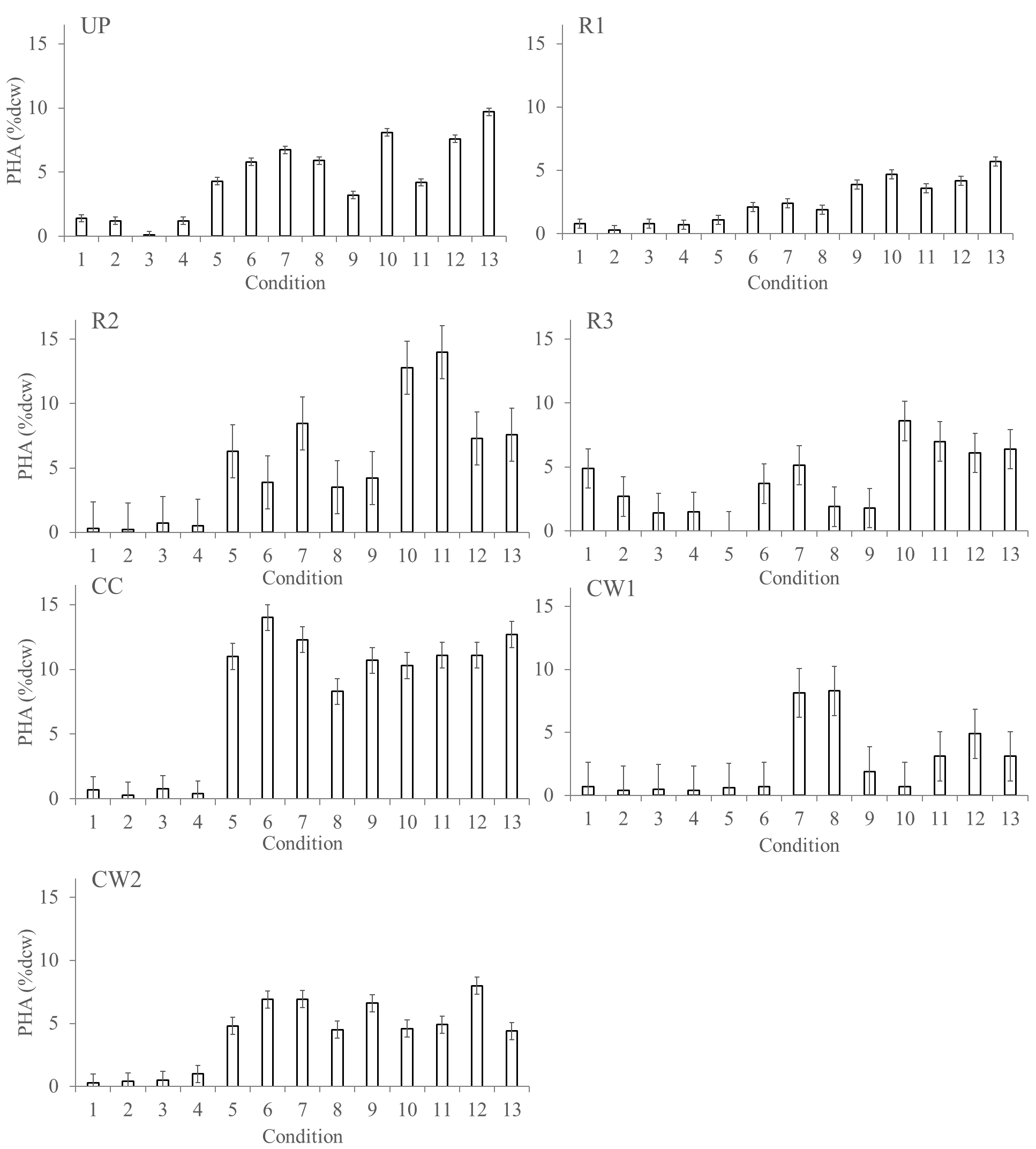

**Figure A2.** PHB production for each microbiome under each of 13 tested conditions (see Table 2 for the properties of conditions).

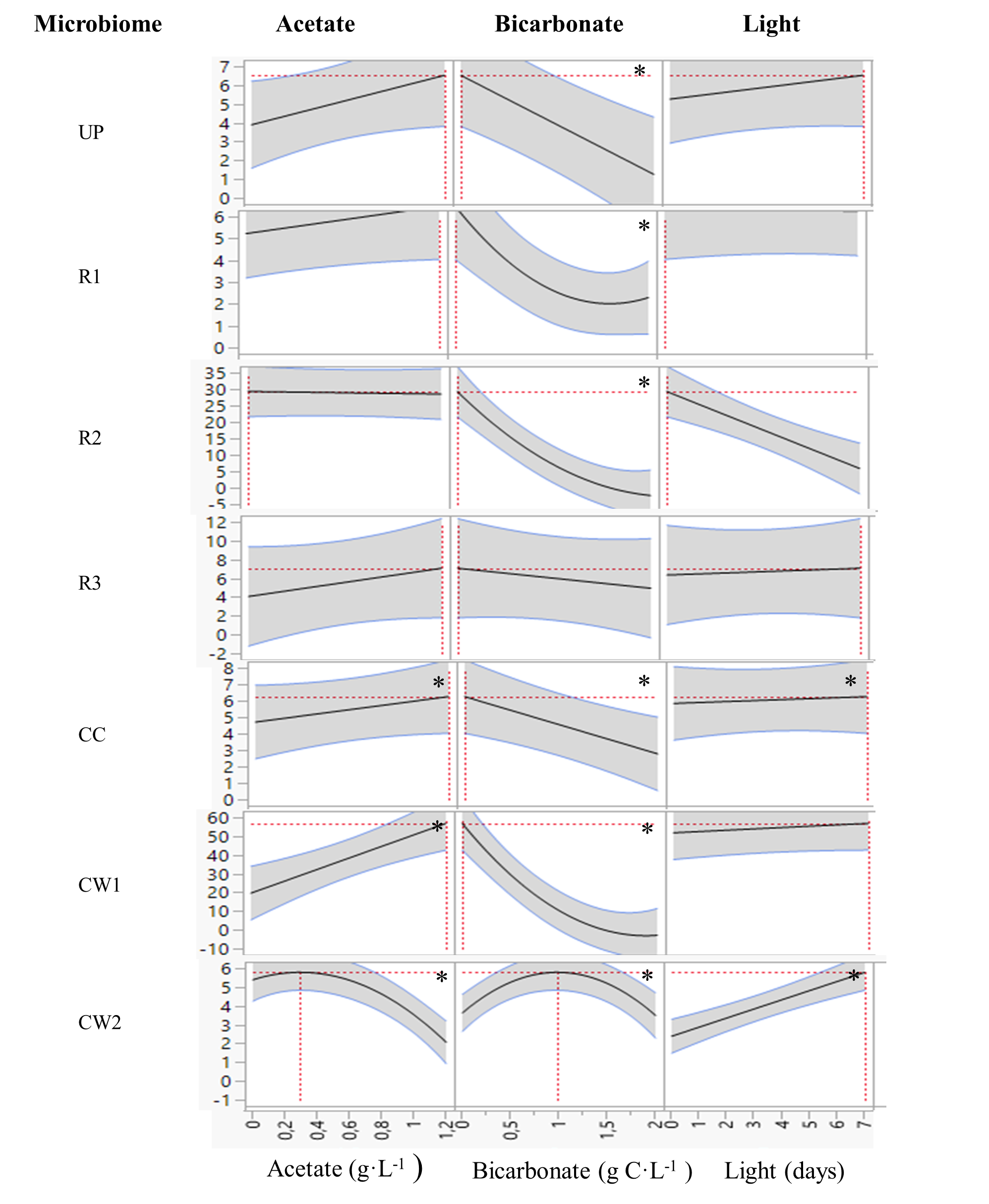

**Figure A3.** Prediction of the effect of acetate, inorganic carbon and light in the EPS production based on DoE results. Illustrating the optimal values of the tested parameters for each microbiome. * indicates parameters which effect is statistically significant. Regression line is shown in black and confidence interval in grey.

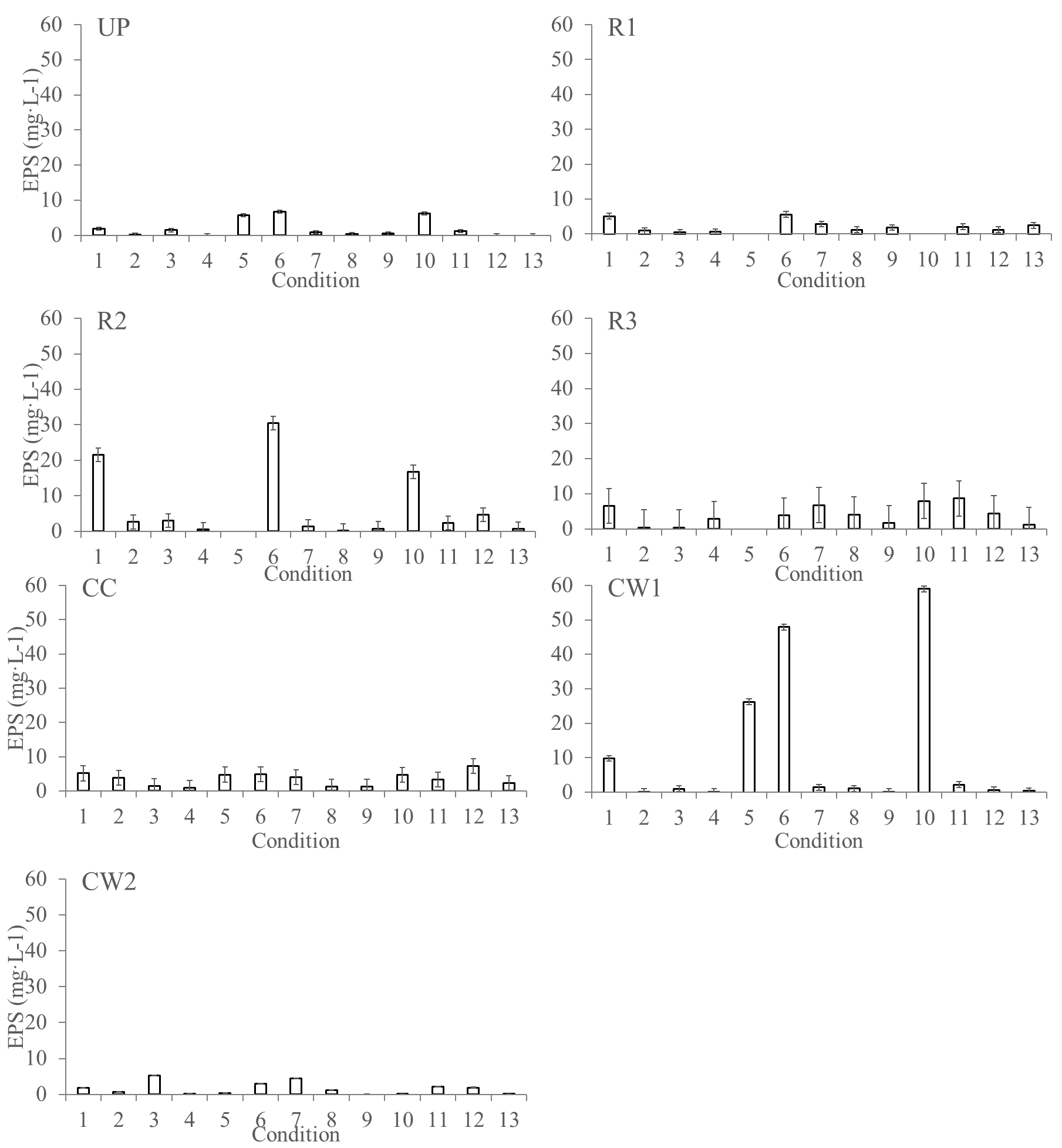

**Figure A4.** EPS production for each microbiome under the 13 tested conditions (see Table 2 for the properties of conditions).

| **Table A2.** Compositional monosaccharides from EPS produced. Data is shown in mg·L^-1^. <lod stands for under detection limit. Lines without number is for absence of mixed liquor before analysis. | | | | | | | | | | |
| --- | --- | --- | --- | --- | --- | --- | --- | --- | --- | --- |
| **Microbiome** | **Condition** | **Fucose** | **Galactose** | **Arabinose** | **Glucose** | **Rhamnose** | **Xylose** | **Mannose** | **Glucuronic acid** | **Total EPS** |
|  | 1 | 0.19 | 0.79 | <lod | 0.30 | 0.24 | 0.15 | <lod | 0.21 | 1.88 |
|  | 2 | <lod | <lod | <lod | 0.03 | <lod | <lod | <lod | 0.07 | 0.10 |
|  | 3 | <lod | 0.10 | <lod | 1.23 | <lod | <lod | <lod | 0.13 | 1.46 |
|  | 4 | <lod | <lod | <lod | <lod | <lod | <lod | <lod | <lod | 0.00 |
|  | 5 | 0.56 | 1.85 | <lod | 0.99 | 1.33 | 0.43 | 0.28 | 0.28 | 5.71 |
|  | 6 | 0.46 | 2.09 | 0.03 | 1.41 | 1.43 | 0.37 | 0.32 | 0.55 | 6.66 |
| UP | 7 | 0.01 | 0.20 | 0.00 | 0.48 | 0.02 | 0.02 | <lod | 0.09 | 0.82 |
|  | 8 | <lod | <lod | <lod | <lod | <lod | <lod | <lod | <lod | 0.00 |
|  | 9 | <lod | 0.05 | <lod | 0.24 | <lod | <lod | <lod | 0.15 | 0.43 |
|  | 10 | 0.32 | 1.99 | <lod | 1.44 | 1.45 | 0.32 | 0.29 | 0.44 | 6.24 |
|  | 11 | 0.02 | 0.31 | <lod | 0.62 | 0.02 | 0.10 | 0.02 | 0.11 | 1.20 |
|  | 12 | <lod | 0.05 | <lod | 0.15 | <lod | <lod | <lod | 0.10 | 0.31 |
|  | 13 | <lod | <lod | <lod | <lod | <lod | <lod | <lod | <lod | 0.00 |
|  | 1 | 0.50 | 1.21 | <lod | 1.15 | <lod | 0.36 | 1.17 | 0.45 | 4.84 |
|  | 2 | 0.08 | 0.11 | <lod | 0.31 | 0.11 | 0.04 | 0.16 | 0.15 | 0.96 |
|  | 3 | 0.02 | <lod | <lod | 0.21 | 0.04 | <lod | 0.07 | 0.11 | 0.44 |
|  | 4 | 0.03 | 0.04 | 0.00 | 0.29 | 0.08 | 0.02 | 0.07 | 0.11 | 0.64 |
|  | 5 |  |  |  |  |  |  |  |  | 0.00 |
|  | 6 | 0.47 | 1.15 | <lod | 0.87 | 1.14 | 0.41 | 0.87 | 0.48 | 5.39 |
| R1 | 7 | 0.19 | 0.27 | <lod | 1.33 | 0.21 | 0.16 | 0.32 | 0.29 | 2.78 |
|  | 8 | 0.02 | 0.00 | <lod | 0.96 | 0.03 | 0.01 | 0.08 | 0.10 | 1.20 |
|  | 9 | 0.06 | 0.08 | <lod | 1.41 | 0.05 | 0.02 | 0.09 | 0.13 | 1.85 |
|  | 10 |  |  |  |  |  |  |  |  | 0.00 |
|  | 11 | 0.20 | 0.19 | <lod | 0.73 | 0.20 | 0.12 | 0.32 | 0.25 | 2.00 |
|  | 12 | 0.03 | 0.04 | <lod | 0.29 | 0.08 | 0.02 | 0.07 | 0.11 | 0.64 |
|  | 13 | 0.08 | <lod | <lod | 2.12 | 0.08 | 0.00 | 0.12 | 0.10 | 2.51 |
|  | 1 | 1.25 | 6.92 | 0.26 | 6.44 | 1.16 | 0.93 | 2.94 | 1.61 | 21.50 |
|  | 2 | 0.24 | 0.98 | <lod | 0.58 | 0.18 | 0.08 | 0.18 | 0.37 | 2.60 |
|  | 3 | 0.15 | 0.91 | <lod | 1.20 | 0.07 | 0.08 | 0.24 | 0.30 | 2.94 |
|  | 4 | <lod | 0.09 | <lod | 0.34 | <lod | <lod | <lod | 0.10 | 0.53 |
|  | 5 |  |  |  |  |  |  |  |  | 0.00 |
|  | 6 | 2.40 | 10.61 | 0.08 | 5.44 | 4.94 | 2.20 | 2.49 | 1.65 | 29.83 |
| R2 | 7 | 0.02 | 0.20 | <lod | 1.02 | <lod | <lod | <lod | 0.13 | 1.37 |
|  | 8 | <lod | 0.00 | <lod | 0.12 | <lod | <lod | <lod | 0.07 | 0.19 |
|  | 9 | <lod | 0.30 | 0.28 | 0.28 | <lod | <lod | <lod | 0.19 | 1.06 |
|  | 10 | 0.82 | 6.42 | 0.16 | 4.57 | 0.92 | 0.55 | 1.47 | 1.25 | 16.16 |
|  | 11 | 0.05 | 1.12 | <lod | 0.80 | 0.06 | <lod | 0.01 | 0.25 | 2.30 |
|  | 12 | 0.33 | 1.79 | <lod | 1.32 | 0.07 | <lod | 0.69 | 0.45 | 4.64 |
|  | 13 | 0.04 | 0.32 | <lod | 0.19 | <lod | <lod | <lod | 0.19 | 0.74 |
|  | 1 | 0.25 | 0.82 | 0.71 | 3.64 | 0.91 | 0.37 | 0.58 | <lod | 7.28 |
|  | 2 | 0.06 | 0.19 | 0.25 | <lod | 0.39 | 0.12 | 0.18 | <lod | 1.18 |
|  | 3 | 0.09 | 0.21 | 0.26 | <lod | 0.38 | 0.13 | 0.19 | <lod | 1.26 |
|  | 4 | 0.07 | 0.51 | 0.45 | 1.75 | 0.46 | 0.21 | 0.29 | 0.01 | 3.76 |
|  | 5 |  |  |  |  |  |  |  |  | 0.00 |
|  | 6 | 0.38 | 0.82 | 0.66 | 1.47 | 0.91 | 0.38 | 0.52 | 0.08 | 5.22 |
| R3 | 7 | 0.24 | 1.28 | 0.72 | 3.20 | 0.86 | 0.45 | 0.64 | 0.14 | 7.52 |
|  | 8 | 0.09 | 0.51 | 0.42 | 3.03 | 0.40 | 0.16 | 0.25 | 0.03 | 4.88 |
|  | 9 | 0.07 | 0.42 | 0.32 | 0.91 | 0.42 | 0.17 | 0.27 | <lod | 2.57 |
|  | 10 | 0.32 | 1.17 | 0.84 | 3.46 | 1.30 | 0.53 | 0.69 | 0.22 | 8.54 |
|  | 11 | 0.29 | 1.62 | 0.99 | 4.30 | 0.99 | 0.59 | 0.74 | 0.17 | 9.70 |
|  | 12 | 0.19 | 0.80 | 0.63 | 2.00 | 0.75 | 0.34 | 0.56 | 0.07 | 5.33 |
|  | 13 | 0.04 | 0.30 | 0.36 | 0.50 | 0.42 | 0.13 | 0.21 | 0.01 | 1.96 |
|  | 1 | 0.65 | 0.98 | 0.47 | 1.16 | 0.26 | 0.17 | 0.35 | 0.47 | 4.50 |
|  | 2 | 0.30 | 0.99 | 0.07 | 1.63 | 0.02 | 0.03 | 0.40 | 0.26 | 3.70 |
|  | 3 | 0.10 | 0.40 | <lod | 0.33 | <lod | <lod | 0.17 | 0.13 | 1.13 |
|  | 4 | 0.05 | 0.17 | <lod | 0.31 | <lod | <lod | 0.03 | 0.08 | 0.66 |
|  | 5 | 0.65 | 1.00 | 0.42 | 0.85 | 0.41 | 0.11 | 0.27 | 0.41 | 4.11 |
|  | 6 | 0.68 | 1.02 | 0.43 | 0.88 | 0.45 | 0.12 | 0.27 | 0.43 | 4.27 |
| CC | 7 | 0.37 | 1.11 | 0.06 | 1.07 | 0.08 | 0.04 | 0.56 | 0.33 | 3.63 |
|  | 8 | 0.15 | 0.38 | <lod | 0.34 | <lod | <lod | 0.15 | 0.15 | 1.17 |
|  | 9 | 0.12 | 0.37 | <lod | 0.49 | <lod | <lod | 0.16 | 0.15 | 1.29 |
|  | 10 | 0.66 | 0.98 | 0.40 | 0.93 | 0.35 | 0.10 | 0.36 | 0.41 | 4.19 |
|  | 11 | 0.30 | 0.92 | <lod | 0.77 | <lod | 0.01 | 0.48 | 0.32 | 2.80 |
|  | 12 | 0.62 | 2.15 | 0.11 | 1.72 | 0.20 | 0.08 | 1.25 | 0.61 | 6.73 |
|  | 13 | 0.26 | 0.69 | <lod | 0.58 | <lod | <lod | 0.36 | 0.23 | 2.12 |
|  | 1 | 1.67 | 0.95 | 0.39 | 3.09 | 1.46 | 0.80 | 0.76 | 0.54 | 9.66 |
|  | 2 | <lod | <lod | <lod | 0.08 | 0.09 | <lod | <lod | <lod | 0.17 |
|  | 3 | 0.12 | 0.13 | <lod | 0.41 | 0.18 | 0.04 | <lod | 0.10 | 0.98 |
|  | 4 | <lod | <lod | <lod | 0.06 | 0.08 | <lod | <lod | <lod | 0.13 |
|  | 5 | 2.81 | 5.65 | 0.67 | 6.47 | 5.05 | 1.76 | 1.79 | 1.67 | 25.86 |
|  | 6 | 6.06 | 10.47 | 0.95 | 13.32 | 8.74 | 3.10 | 2.88 | 1.94 | 47.46 |
| CW1 | 7 | 0.19 | 0.37 | 0.01 | 0.50 | 0.11 | 0.09 | 0.04 | 0.12 | 1.43 |
|  | 8 | 0.15 | 0.20 | <lod | 0.36 | 0.19 | 0.04 | <lod | 0.10 | 1.04 |
|  | 9 | <lod | <lod | <lod | 0.05 | 0.08 | <lod | <lod | <lod | 0.12 |
|  | 10 | 5.55 | 13.37 | 1.31 | 19.18 | 9.77 | 3.12 | 3.28 | 2.76 | 58.35 |
|  | 11 | 0.01 | 0.14 | <lod | 1.80 | 0.17 | 0.02 | <lod | 0.08 | 2.22 |
|  | 12 | 0.02 | 0.10 | <lod | 0.33 | 0.12 | <lod | <lod | 0.07 | 0.64 |
|  | 13 | <lod | 0.06 | <lod | 0.17 | 0.11 | <lod | <lod | 0.07 | 0.41 |
|  | 1 | 0.01 | 0.09 | 0.45 | <lod | <lod | <lod | <lod | 0.11 | 0.66 |
|  | 2 | <lod | <lod | 0.11 | <lod | <lod | <lod | 0.06 | 0.14 | 0.30 |
|  | 3 | 0.43 | 1.62 | <lod | 1.65 | <lod | 0.09 | 1.30 | 0.16 | 5.24 |
|  | 4 | <lod | <lod | 0.04 | <lod | <lod | <lod | <lod | <lod | 0.04 |
|  | 5 | <lod | <lod | 0.18 | <lod | <lod | 0.01 | 0.05 | 0.10 | 0.34 |
|  | 6 | 0.20 | 0.62 | <lod | 1.09 | <lod | 0.06 | 0.93 | 0.11 | 3.02 |
| CW2 | 7 | 0.35 | 0.58 | 0.61 | 1.49 | <lod | 0.14 | 1.06 | 0.14 | 4.35 |
|  | 8 | <lod | 0.24 | 0.27 | <lod | <lod | <lod | 0.12 | 0.44 | 1.07 |
|  | 9 | <lod | <lod | <lod | <lod | <lod | <lod | <lod | <lod | 0.00 |
|  | 10 | <lod | <lod | <lod | <lod | <lod | <lod | 0.01 | <lod | 0.01 |
|  | 11 | <lod | 0.02 | <lod | <lod | <lod | 0.01 | 0.07 | 0.07 | 0.17 |
|  | 12 | 0.40 | 1.25 | <lod | 1.73 | <lod | 0.22 | 0.22 | 0.15 | 3.98 |
|  | 13 | <lod | <lod | <lod | <lod | <lod | <lod | <lod | 0.09 | 0.09 |
